# Increased slow wave activity in response to sleep deprivation is highest immediately after micro-arousals

**DOI:** 10.1101/2025.01.29.635246

**Authors:** Natalie L. Hauglund, Lukas B. Krone, Martin Kahn, Cristina Blanco-Duque, Vladyslav V. Vyazovskiy

## Abstract

Non-rapid eye movement (NREM) sleep is not uniform but characterized by brief intrusions of wake-like brain activity, known as micro-arousals or brief awakenings. Although micro-arousals are an inherent feature of both human and animal sleep, knowledge about the neural correlates of micro-arousals across the cortical column is sparse. We here developed an algorithm for automatic detection of micro-arousals based on EMG activity and used it to explore how micro-arousals modulate laminar neural activity in the motor cortex, and how this relationship is affected by sleep pressure in mice. Our analysis showed that micro-arousals were associated with a general suppression of local field potential (LFP) power across the cortical column. Slow wave activity (SWA, 1-4 Hz) immediately after micro-arousals was tightly correlated with sleep pressure and even surpassed average SWA levels during NREM sleep in sleep deprived animals. In addition, analysis of single-channel firing showed that some channels exhibited increased activity immediately prior to micro-arousals, while others exhibited decreased activity. This study provides new insights into the neural mechanisms underlying micro-arousals and identifies a new link between micro-arousals and sleep homeostasis.

## Introduction

Sleep is a prerequisite for health, whereas sleep disruption is associated with a broad range of pathological conditions. Healthy and restorative sleep rests on the ability to maintain long and uninterrupted periods of sleep. However, non-rapid eye movement (NREM) sleep is interspersed by regular arousals with a brief shift of the electroencephalogram (EEG) to more wake-like frequencies followed by transition directly back to NREM sleep. These micro-arousals, or brief awakenings, are normally not consciously perceived, occur even in the absence of external stimuli, and their functional significance is not known^1,2^.

Micro-arousals result from internally-generated infraslow oscillations in noradrenaline (NA) released from the brain stem nucleus locus coeruleus (LC) during NREM sleep^3–5^. A wide range of theories on the function of micro-arousals have been proposed. Some suggest that they have a homeostatic function to maintain or restore physiological processes, such as preventing that sleep becomes too deep^6^, preventing premature entry to REM sleep^6^, or to restore cardiorespiratory function in sleep^7^. Another theory is that micro-arousals provide an opportunity to scan the environment for potential dangers without disrupting sleep. Recent studies showed that infraslow cycles in activity of the LC during sleep is implicated in memory formation^4^ and drives cerebrospinal fluid flow, and thereby cleaning of the brain via the glymphatic system during sleep^8^. Thus, the mechanisms responsible for generating micro-arousals have a crucial role in the restorative processes that take place during sleep^2^.

The neuronal activity pattern associated with micro-arousals is distinct from wakefulness^9^, indicating that micro-arousals are not the same as actual awakenings. The number of arousals from sleep exhibit a strong negative correlation with self-reported sleep quality^10^. However, many studies do not clearly distinguish between full awakenings and micro-arousals, which makes it difficult to assess the association between internally-generated micro-arousals and subjective sleep quality or pathology. Factors such as stress have been shown to increase the quantity of micro-arousals due to an increase in the frequency of NA oscillations^5,11^. Similarly, old age is associated with an increase in micro-arousals^12–14^, suggesting that an elevated micro-arousal frequency is indicative of poor sleep. This association between micro-arousals and poor sleep quality has contributed to the common misconception that all micro-arousals *per se* are equivalent to sleep fragmentation and poor sleep. The relationship between sleep arousals and all-cause mortality is a bell-shaped curve where both a very low number and a high number of micro-arousals are associated with higher mortality rate^15^. Thus, the relationship between micro-arousals and sleep quality is more complicated than “the fewer the better”, which underscores the importance of gaining a better understanding of the interplay between micro-arousals and sleep physiology.

Sleep is under homeostatic regulation and the amplitude of EEG slow wave activity (SWA, typically defined as EEG oscillations between 0.5-4 Hz) during NREM sleep increases as a function of time spent awake prior to the sleep episode^16–18^. The generation of micro-arousals is also homeostatically regulated as high sleep pressure is linked to fewer micro-arousals in both humans^19^ and rodents^20,21^. However, it is currently not known whether neuronal activity associated with micro-arousals is affected by sleep pressure.

Manual scoring of micro-arousals is time consuming and can be affected by subjectivity. Furthermore, micro-arousals are brief events with a duration that often falls below four seconds, which is the bin size typically used for rodent sleep scoring. This makes accurate analysis of neuronal activity around micro-arousals difficult. To overcome these challenges, we here developed an algorithm to automatically detect NREM sleep micro-arousals based on the electromyography (EMG) signal. We subsequently used the algorithm to detect micro-arousals in continuous sleep data from a group of mice with laminar local field potential (LFP) and multiunit activity (MUA) recordings from the motor cortex. Recordings from the mice during 24 hours of baseline as well as after 6 hours of sleep deprivation allowed us to investigate cortical neuronal activity across the motor cortex during micro-arousals, and assess how these activity patterns are impacted by sleep pressure.

## Results

### Automated algorithm for detection of NREM micro-arousals is a useful tool for objective micro-arousal detection

In order to precisely assess the neural correlates of micro-arousals, we aimed to detect micro-arousals and their start and end times with high temporal accuracy. We therefore developed an algorithm that detects micro-arousals during NREM sleep based on analysis of the EMG and predefined selection criteria. While several definitions of micro-arousals exist^4,5,8,22^, we here defined micro-arousals as brief (below 5 seconds) bouts of muscle activity during NREM sleep. The algorithm was developed to work as a post-processing tool on manually-scored sleep data that had already been segmented into the three main vigilance states: wakefulness, NREM sleep and REM sleep. This allowed us to use the algorithm on already obtained datasets in order to re-analyse them with a focus on micro-arousals. Only the EMG signal was used for the detection to allow investigation of the neuronal dynamics in an unbiased manner. The processing pipeline for the algorithm is shown in Figure 1A. After inputting the EMG trace and the corresponding vigilance state scoring, the EMG was pre-processed and EMG activity peaks were included or excluded based on criteria such as duration and amplitude (see detailed description in method section). The resulting EMG peaks represented NREM micro-arousals that were used for analysis. An average of the rectified EMG signal during the detected micro-arousals confirmed that the algorithm successfully identified brief increases in EMG tonus (Fig. 1B).

**Figure 1.**
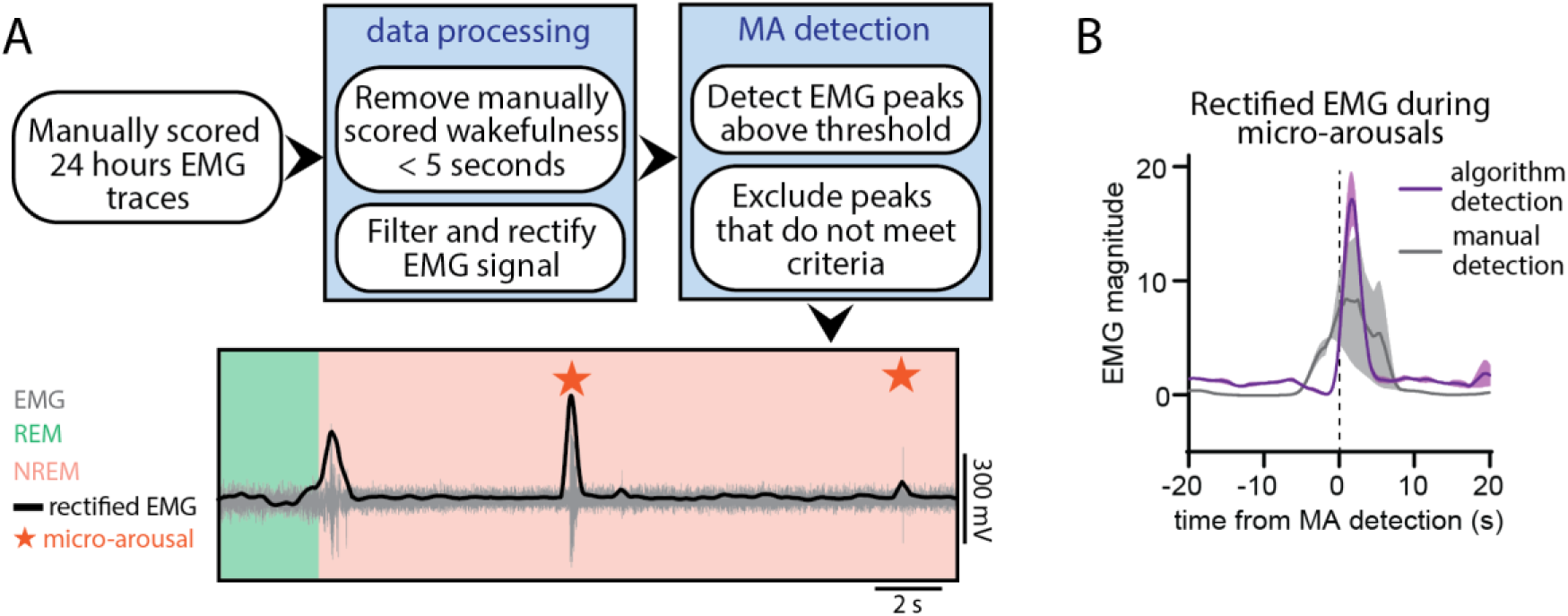
Automated algorithm for detection of NREM micro-arousals is a useful tool for objective micro-arousal detection. (A) The sequential steps of the pipeline for automated detection of micro-arousals. The pipeline is used on EMG data pre-scored for vigilance states. In the first steps, manually scored states with a duration below 10 seconds between two NREM episodes are removed from the scoring to omit subjectively scored micro-arousals. The EMG signal is then filtered and rectified, and EMG peaks during NREM sleep are detected. During the next steps, EMG peaks that do not occur during NREM sleep are excluded and peaks immediately following each other are combined. Finally, EMG peaks that follows REM sleep and EMG peaks with a duration above 5 seconds are excluded. The last image is an example of the resulting scoring showing the original EMG signal in grey, the rectified EMG used for peak detection in black and the detected micro-arousals as orange stars. Manually scored behavioural states showed as shading in the background. Notice how micro-arousals following REM sleep are not included. (B) Mean rectified EMG trace during micro-arousals (MAs) in comparison with algorithm-based and manual detection (n=7, shaded area is SEM).

### Micro-arousals elicit a decrease in LFP power throughout layers of the motor cortex

The micro-arousal detection algorithm was applied to a dataset previously published^23–25^. In this dataset, adult wild-type (C57BL/6) mice were implanted with electrodes for recording laminar LFP and MUA from the left primary motor cortex (M1), concomitantly with EEG and EMG recordings. We first set out to investigate the neuronal activity pattern during micro-arousals across cortical layers (Fig. 2A). 24 hours of baseline data was manually scored as wakefulness, NREM sleep and REM sleep based on the EEG and EMG signal (Fig. 2B), and LFP and MUA was extracted for analysis (Fig. 2C). Spectral analysis revealed a brief but consistent drop in LFP power across layers 2/3, 4, 5, and 6 at the onset of micro-arousal initiation followed by a gradual increase back to baseline (Fig. 2D). The drop in power was evident for all power bands between 1-30 Hz while gamma power (30-80 Hz) exhibited a brief peak during the micro-arousal followed by a slow increase (Fig. 2E). Power spectral densities 30-50 seconds before the micro arousal compared to during the micro-arousal (0-2 seconds after micro-arousal onset) confirmed a drop in power in all M1 layers (Fig. 2F). Quantification of the power decrease from before to during micro-arousals revealed that all layers displayed a ∼50-70% decrease in power and that the most pronounced drop happened in the slow wave activity (SWA) / delta (1-4 Hz) frequency range (Fig. 2G). Although this was true for all layers, layer 5 exhibited a slightly larger drop compared to layer 2/3 (Fig. 2H). Overall, the analysis showed that micro-arousals were associated with a pronounced decrease in LFP power across frequencies and cortical layers.

**Figure 2.**
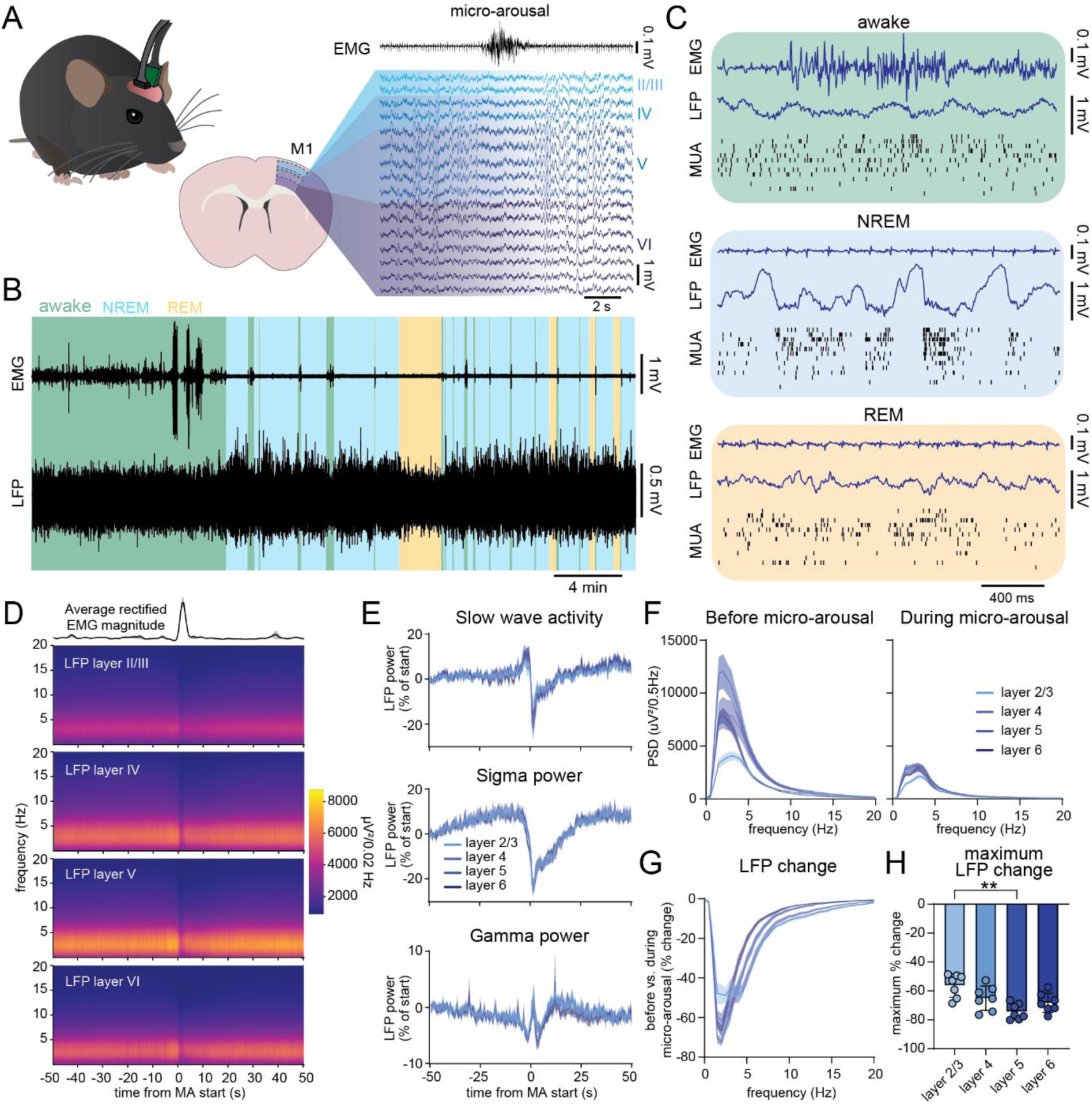
Micro-arousals elicit a decrease in LFP power throughout layers of the motor cortex. (A) Diagram of the experimental setup. EMG, EEG and LFP traces from the motor cortex were acquired from C57Bl/6 mice. Micro-arousals were detected by the algorithm and the LFP traces around every micro-arousal were analysed. (B) Example EMG and LFP traces with the background colour coded to show vigilance states. (C) Example EMG, LFP and multi-unit activity data from wakefulness, NREM sleep and REM sleep. (D) Top: mean rectified EMG trace and bottom: mean LFP spectrogram from layer 2/3, layer 4, layer 5, and layer 6. All are time-locked to the beginning of the micro-arousal start (n=7). (E) SWA, sigma and gamma power across layers 2/3, 4, 5, and 6 time-locked to the beginning of micro-arousals. Power is calculated as the percent change from 30-50 s before (n=7). (F) Power spectral densities for layers 2/3, 4, 5, and 6 before (t=-50 to-30 s) and during (t=0 to 2 s) micro-arousals (n=7). (G) Quantification of the percentage change between before and during micro-arousals for all layers (n=7). (H) Quantification of the maximum change from data shown in (G) (n=7, RM one-way ANOVA with Geisser-Greenhouse correction and Tukey’s multiple comparison). **P=<0.01. Shaded area around line graphs are ±SEM.

### Post micro-arousal SWA exceeds NREM sleep SWA in sleep deprived mice

Sleep homeostasis refers to the ability to keep track of time spent in sleep and wakefulness, and ensures that the need for an adequate amount of sleep is satisfied. High sleep drive manifests as increased SWA in NREM sleep and an increased threshold to be woken up by external stimuli^17,26–28^. We next set out to investigate if sleep homeostasis affects the neuronal characteristics of micro-arousals by analysing neuronal firing patterns during micro-arousals under conditions of high or low sleep pressure (Fig. 3A). Micro-arousals were pooled into three groups; 1) low sleep pressure (micro-arousals occurring during the last 6 hours of the light phase), 2) medium sleep pressure (micro-arousals occurring during the first 6 hours of the light phase) and 3) high sleep pressure (micro-arousals occurring during the first 2 hours of recovery sleep after 6 hours of sleep deprivation by novel objects exposure started at the beginning of the light phase). Neuronal activity around micro-arousals was then compared between the groups. Spectral analysis of layer 5 motor cortex LFP signals confirmed that higher sleep pressure was associated with increased levels of NREM SWA (Fig. 3B). The amplitude of the EMG signal during micro-arousals exhibited a larger variance in the high sleep pressure group, but no significant difference was observed between groups (Fig. 3C). Analysis of LFP power during micro-arousals revealed that all three groups exhibited an increase in SWA right before a micro-arousal, but SWA immediately after the micro-arousal was highly dependent on sleep pressure (Fig. 3D-E). The low sleep pressure group exhibited a long period with decreased SWA following micro-arousals, while the high sleep pressure group exhibited a pronounced SWA increase with SWA levels exceeding those before the micro-arousal. Power spectral density analysis of the period before (30-50 seconds before muscle activity), during (0-2 seconds after the first muscle activity), and after (5-10 seconds after the end of the muscle activity) each micro-arousal showed that all groups exhibited a decrease in power during micro-arousals (Fig. 3F). In addition, the analysis confirmed that the low and medium sleep pressure groups showed reduced SWA levels after micro-arousals, while the high sleep pressure group showed higher SWA levels. The percentage change in power spectral densities before versus after the micro-arousal was highly sensitive to sleep/wake history (Fig. 3G). Both the low and medium sleep pressure group exhibited a decrease in low frequencies after compared to before the micro-arousal. The low sleep pressure group displayed significantly larger negative changes than the medium sleep pressure group in frequencies between 0.5-6 Hz. Contrary to the low and middle sleep pressure group, the high sleep pressure group displayed a significant positive change in power spectral densities from before to after the micro-arousal. The difference between low and high sleep pressure spanned a frequency range from 0-7.5 Hz. This sleep-pressure sensitive shift was also evident when the percentage change was plotted as the area under the curve (Fig. 3H). Analysis of the duration of micro-arousals during a 24-hour baseline recording compared with a period 2 hours after sleep deprivation showed that this was not due to differences in the duration of micro-arousal between low and high sleep pressure conditions (Fig. S1A).

**Figure 3.**
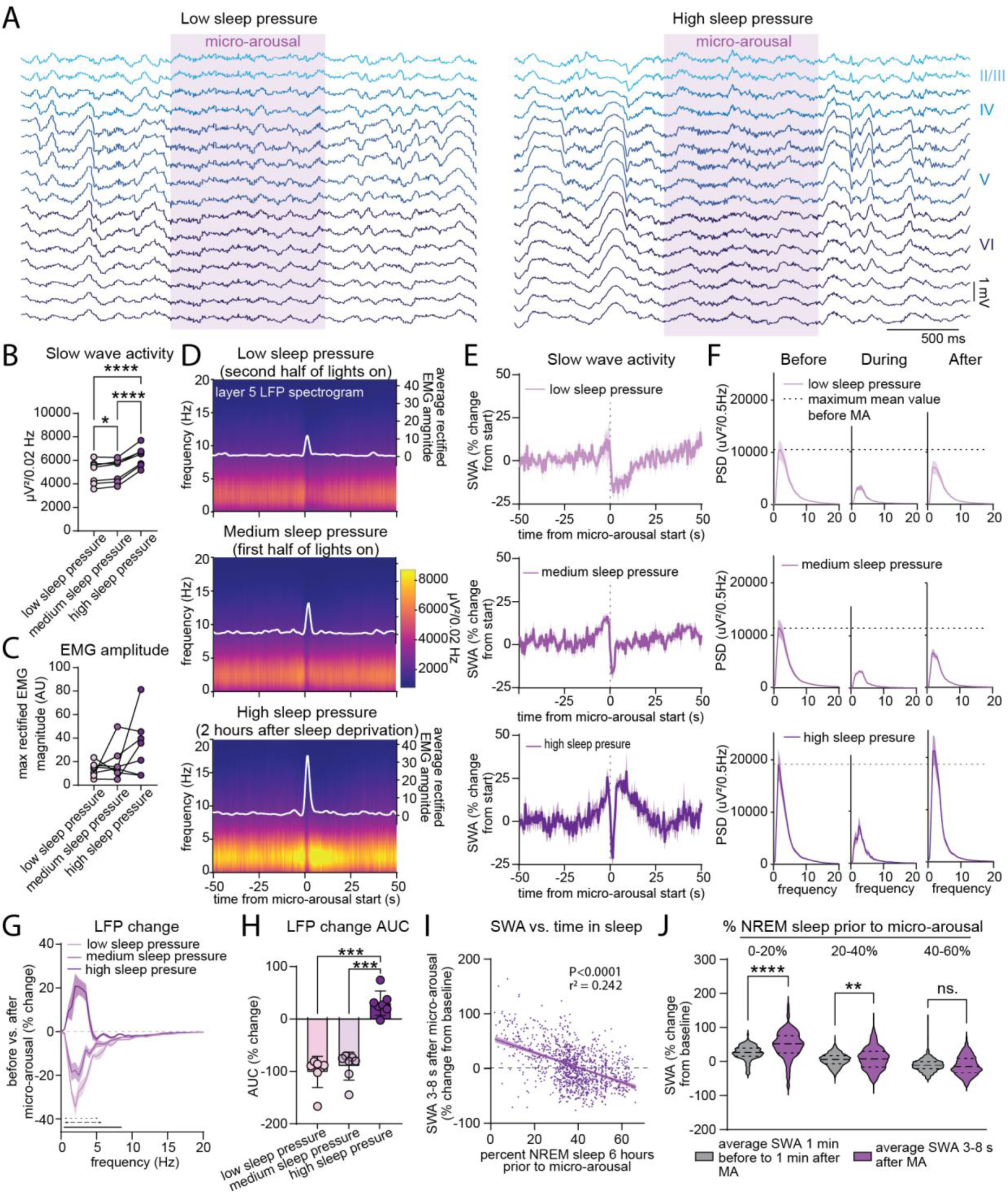
Post micro-arousal SWA exceeds NREM sleep SWA in sleep deprived mice. (A) Example layer 5 motor cortex LFP traces during a micro-arousal under baseline conditions (low sleep pressure, right) and after 6 hours of sleep deprivation (high sleep pressure, left). (B-C) NREM SWA (B) and amplitude of EMG activity during micro-arousals (C) within the low sleep pressure time bin (last 6 hours of the light phase), medium sleep pressure time bin (first 6 hours of the light phase), and high sleep pressure time bin (first 2 hours after 6 hours sleep deprivation) (n=7, RM one-way ANOVA with Geisser-Greenhouse correction and Tukey’s multiple comparison). (D) Mean LFP spectrograms from layer 5 neurons during low, medium, and high sleep pressure where t=0 depicts start of EMG activity (n=7). (E) SWA expressed as percent change from 50-20 s before the micro-arousal during low (top panel), medium (middle panel) or high (bottom panel) sleep pressure (n=7, shaded area is ±SEM). (F) Mean power spectral density plots before (t=-50 to-30 s), during (t=0 to 2 s) and after (t=5 to 10 s) micro-arousals occurring under low (top panel), medium (middle panel) or high (bottom panel) sleep pressure (n=7, shaded area is ±SEM, dashed horizontal line shows the maximum value before the micro-arousal). (G) Percentage change in power spectral densities before to after the micro-arousal during low, medium and high sleep pressure. Black line denotes significant difference between medium and low sleep pressure (0.5-6 Hz), black dashed line denotes significant different between low and high sleep pressure (0-7.5 Hz) and black dotted line denotes significant difference between medium and high sleep pressure (0-4.5 Hz) (n=7, RM two-way ANOVA and Tukey’s multiple comparison, shaded area is ±SEM). (H) Quantification of mean area under the curve for data in (G) (n=7, error bars are SD, RM one-way ANOVA with Geisser-Greenhouse correction and Tukey’s multiple comparison). (I) Percent time spent in NREM sleep during a period 6 hours before each micro-arousal correlated with the amplitude of SWA 3-8 seconds after the same micro-arousal. SWA amplitude is shown as percent of mean NREM SWA during the first 6 hours of the light phase on the baseline day (Pearson correlation, 1162 micro-arousals from 7 mice during 18 hours after 6 hours sleep deprivation included, the line shows the linear regression and the shaded area depict 95% confidence intervals). (J) Comparison of average SWA (shown as the percent change from NREM SWA during the first 6 hours of the light phase on the baseline day) during a period 1 minute before to 1 minute after each micro-arousal and during a period 3-8 seconds after each micro-arousal. The micro-arousals are divided into bins after the amount of NREM sleep that was present 6 hours before the micro-arousal (n=7 with 116 micro-arousals in the 0-20% bin, 529 micro-arousals in the 20-40% bin, and 457 micro-arousals in the 40-60% bin, Paired t-test, P<0.0001 and P=0.001). Shaded area around line graphs are ±SEM. *P=<0.05, **P=<0.01, ***P=<0.001, ****P=<0.0001, MA=micro-arousal, SWA=SWA, LFP=local field potential.

It is well-established that NREM sleep SWA reflects time spent awake in the period prior to sleep^17,18,27^. We next asked if the period immediately following micro-arousals follow the same dynamics. First, all micro-arousals during a period of 18 hours following 6 hours sleep deprivation with novel objects (1162 micro-arousals from 7 mice) were collected for analysis. The level of SWA 3-8 seconds after each micro-arousal was then plotted against the percent time spent in NREM sleep during 6-hours prior to the micro-arousal (Fig. 3I). Post-arousal SWA exhibited a significant negative correlation with prior time spent in sleep, meaning that less sleep resulted in higher SWA, in accordance with the established link between SWA and sleep homeostasis. To investigate the relationship between SWA during NREM sleep compared to post-arousal SWA, we calculated SWA during a 2-minute period around each micro-arousal (1 minute before to 1 minute after). Post-arousal SWA (3-8 seconds after the micro-arousal) exhibited a significant positive correlation with SWA during the 2-minute period around the micro-arousal (Fig. S1B). However, the slope of the correlation between sleep time and post-arousal SWA was significantly steeper than the slope of the correlation between sleep time and SWA 1 minute before to 1 minute after the micro-arousal (Fig. S1C). This indicates that post-arousal SWA is more sensitive to homeostatic sleep pressure than general NREM sleep SWA. We found that post-arousal SWA was significantly higher than SWA 1 minute before to 1 minute after the micro-arousal when the mice had only spent 0-20% and 20-40% of the past 6 hours in sleep but not if they had spent 40-60% in sleep (Fig. 3J). In addition, this effect was stronger for mice in the 0-20% group than the 20-40% group, indicating that the difference between post-arousal SWA and NREM SWA is more pronounced with increasing levels of sleep pressure. Overall, these results identifies a short time window around the transition from micro-arousals back to NREM sleep that is especially sensitive to sleep/wake history.

### Neuronal activity during micro-arousals following REM sleep differ from NREM sleep micro-arousals

Micro-arousals occur regularly during NREM sleep but are also commonly observed after bouts of REM sleep upon the transition back to NREM sleep. Micro-arousals during NREM sleep and micro-arousals at the termination of REM sleep both occur alongside an increase in locus coeruleus activity and resulting increase in noradrenaline release^4^. However, how similar the two types of micro-arousals are in terms of neuronal dynamics and sensitivity to sleep homeostasis is not well established. We used an altered version of the micro-arousal detection algorithm to only detect micro-arousals that followed REM sleep and analysed the neuronal activity patterns. Slow waves were regularly observed in the layers of the motor cortex during REM sleep, as previously described^29^ (Fig. 4A). Spectral analysis of micro-arousals at the termination of REM sleep showed a gradual decrease in SWA before the micro-arousal, which was followed by a gradual increase in SWA after the transition to NREM sleep (Fig. 4B). REM sleep micro-arousals exhibited significantly lower SWA levels than NREM sleep micro-arousals (Fig. 4C). Finally, there was an overall significant difference between post-micro-arousal (5-10 seconds after micro-arousal initiation) SWA under different sleep pressure conditions (first 6 hours of lights on, last 6 hours of lights on, first 2 hours after 6 hours sleep deprivation) but no significant difference between the individual sleep pressure conditions in post-hoc analysis (Fig. 4D).

**Figure 4.**
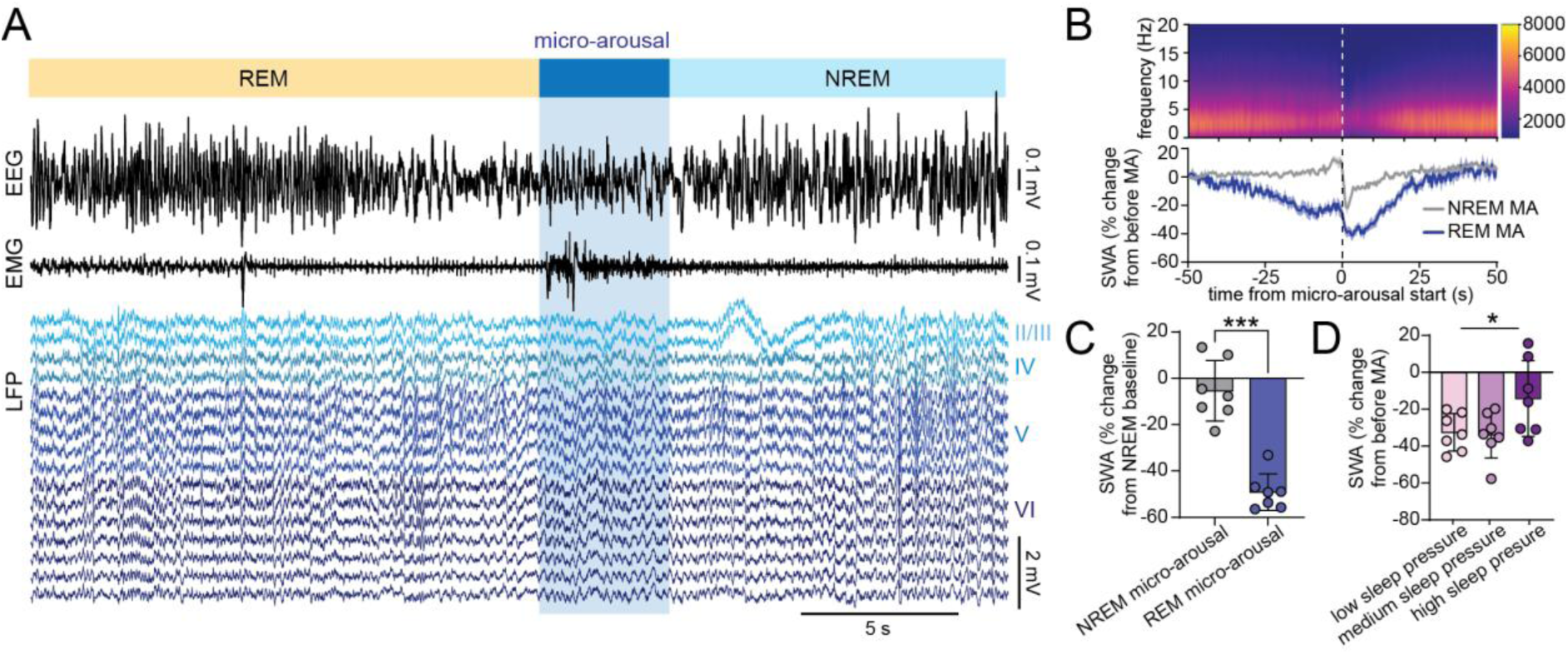
Neuronal activity during micro-arousals following REM sleep differ from NREM sleep micro-arousals. (A) Example of EEG, EMG, and motor cortex LFP traces showing REM sleep termination followed by a micro-arousal and transition to NREM sleep. (B) LFP spectrogram (top) and mean SWA expressed as percent change from 50-20 s before the micro-arousal (bottom) with t=0 depicting the beginning of the micro-arousal (n=7, shaded area is ±SEM). (C) Mean SWA 0-5 seconds after onset of NREM and REM micro-arousals as percent of NREM baseline SWA (n=7, all micro-arousals during the 24 hour baseline recording included, error bars are SD). (D) Mean change in SWA from 50-20 s before to 5-10 seconds after REM sleep-terminating micro-arousals during low, medium and high sleep pressure (n=7, error bars are SD, RM one-way ANOVA with Geisser-Greenhouse correction and Tukey’s multiple comparison, Repeated measures effect P=0.0435, no significant effect in the multiple comparisons test). MA=micro-arousal, SWA=SWA, LFP=local field potential.

### Multi-unit activity recordings from cortical neurons display opposing firing patterns during micro-arousals

In order to better understand the neuronal dynamics of micro-arousals, we next conducted an analysis of single LFP channels across the motor cortex layers. Calculation of the cumulated firing rate across LFP channels showed that micro-arousals with a duration below 5 seconds were associated with a temporary decrease in firing rates that reached the lowest level right after the first muscle activity was detected (Fig. 5A-B). Interestingly, the average firing rate pattern exhibited a short period of briefly increased activity immediately prior to the beginning of the micro-arousal. This “shoulder” of increased firing before was barely visible during low sleep pressure conditions but was significantly increased during the high sleep pressure condition (Fig 5C-D). On the contrary, the decrease in firing did not show significant differences between sleep pressure conditions when all channels were averaged. To investigate if the dual response in average firing rate was due to different firing patterns across channels, all channels were sorted based on whether they displayed increased or decreased firing during micro-arousals under the high sleep pressure condition. Channels showing an increase or a decrease larger than 5 times the standard deviation of the firing rate during the first 15 seconds of the trace, were collected for analysis. A total of 9 channels from 4 different mice were found to show increased activity during micro-arousals (Fig. 5F). Their micro-arousal associated activity did not differ between low and medium sleep pressure conditions, but was significantly increased under high sleep pressure (Fig. 5G) with a peak 0.67 ±0.47 s before the beginning of the micro-arousal. A total of 12 channels from 6 different mice were found to show decreased firing during micro-arousals (Fig. 5H). These channels also exhibited sensitivity to sleep pressure, although the smallest decrease in firing rates were found in the medium sleep pressure group and the most pronounced drop was evident under high sleep pressure (Fig. 5I) with lowest values 1.33 ±2.95 s after the initiation of the micro-arousal. Channels with decreased and increased firing during micro-arousals were scattered across the cortical column (Fig. S2). In summary, analysis of neuronal firing during micro-arousals reveals a non-uniform pattern where some channels exhibit increased firing immediately prior to the micro-arousal while other channels exhibit decreased firing that reaches a minimum immediately after the first muscle activity has been detected.

**Figure 5.**
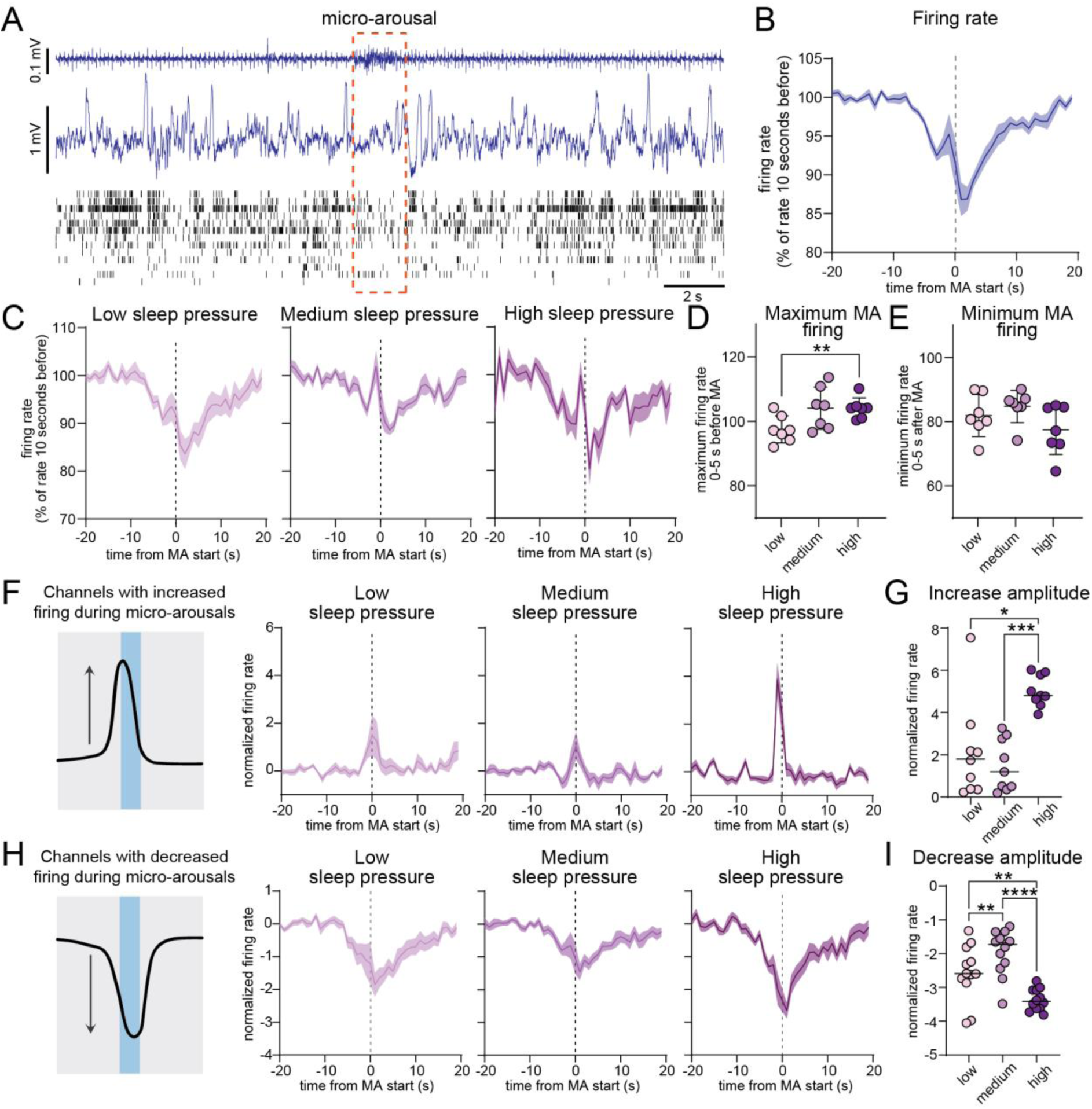
Multi-unit activity recordings from cortical neurons display opposing firing patterns during micro-arousals. (A) Representative EMG, LFP and multi-unit activity from the motor cortex during a micro-arousal. (B) Average firing rate as number of spikes per second time locked to the beginning of micro-arousals (n=7). (C) Firing rate during micro-arousals while mice were subjected to low, medium and high sleep pressure (n=7). (D) Quantification of maximal firing rates in the period 5 seconds before the beginning of micro-arousals (n=7, RM one-way ANOVA with Geisser-Greenhouse correction and Tukey’s multiple comparison). (E) Quantification of minimum firing rates in the period 5 seconds after the beginning of micro-arousals (n=7, RM one-way ANOVA with Geisser-Greenhouse correction and Tukey’s multiple comparison). (F) Average firing rate under different sleep pressure conditions for channels exhibiting increased firing during the high sleep pressure condition (total of 9 channels from 4 mice, values are normalized to the standard deviation). (G) Amplitude of the peak in firing rates from 5 s before to 5 s after the beginning of micro-arousals arousals (n=7, RM one-way ANOVA with Geisser-Greenhouse correction and Tukey’s multiple comparison). (H) Average firing rate under increasing sleep pressure for channels exhibiting decreased firing during low sleep pressure condition (total of 12 channels from 6 mice, values are normalized to the standard deviation). (I) Amplitude of the minimum firing rate from 5 s before to 5 s after the beginning of micro-arousals arousals (n=7, RM one-way ANOVA with Geisser-Greenhouse correction and Tukey’s multiple comparison). Shaded area around line graphs are ±SEM), MA=micro-arousal.

### The duration of micro-arousals affects cortical firing

From the raw multi-unit activity traces, we noted that not all micro-arousals were associated with decreased firing, but some seemed to instead coincide with increased firing compared to the surrounding NREM sleep (Fig. 6A). Analysis of the durations of micro-arousals across mice showed that most micro-arousals lasted around 3 seconds while only 30% of micro-arousals were between 5 and 10 seconds (Fig. 6B). We hypothesized that short micro-arousals were associated with a decrease in firing while longer micro-arousals would have more similarity to wakefulness that generally have higher firing rates than NREM sleep^30^. To test this hypothesis, we increased the upper limit for micro-arousal duration to 10 seconds and ran the detection algorithm. We then divided micro-arousals into groups based on their duration so one group consisted of micro-arousals with a duration below 3 seconds and the other group consisted of micro-arousals with a duration from 5 to 10 seconds. To get a sufficient separation between shorter and longer lasting micro-arousals for this analysis, micro-arousals with a duration of 4 seconds were not included. Comparison of the average firing rate, measured as number of spikes per second, showed significantly higher values for micro-arousals with a duration between 5 and 10 seconds (Fig. 6C-D). To test if this difference was due to increased firing within micro-arousal-active channels or a switch in micro-arousal-inactive channels from silent to active, we collected channels where micro-arousals with a duration below 3 seconds were associated with either an increase or decrease larger than 2 times the standard deviation during the first 15 seconds of the trace. A lower threshold was used here than in the previous analysis to be able to include a sufficient number of micro-arousals in each group. Channels with increased firing did not show a significant difference in firing between short and longer micro-arousals (Fig. 6E-F). However, channels exhibiting decreased firing during short micro-arousals had significantly higher firing during longer micro-arousals (Fig. 6G-H). Because the resolution of the multi-unit activity probes does not allow for precise identification of single neurons, it is not possible to say whether the increased firing is the result of a shift in activity of specific neurons from silent to active, or whether it reflects that longer lasting micro-arousals engage a larger number of neurons in proximity to the probe. However, these results indicate that micro-arousals during NREM sleep are not uniform and are associated with a complex interplay between different groups of neurons that is affected by the duration of the micro-arousal.

**Figure 6.**
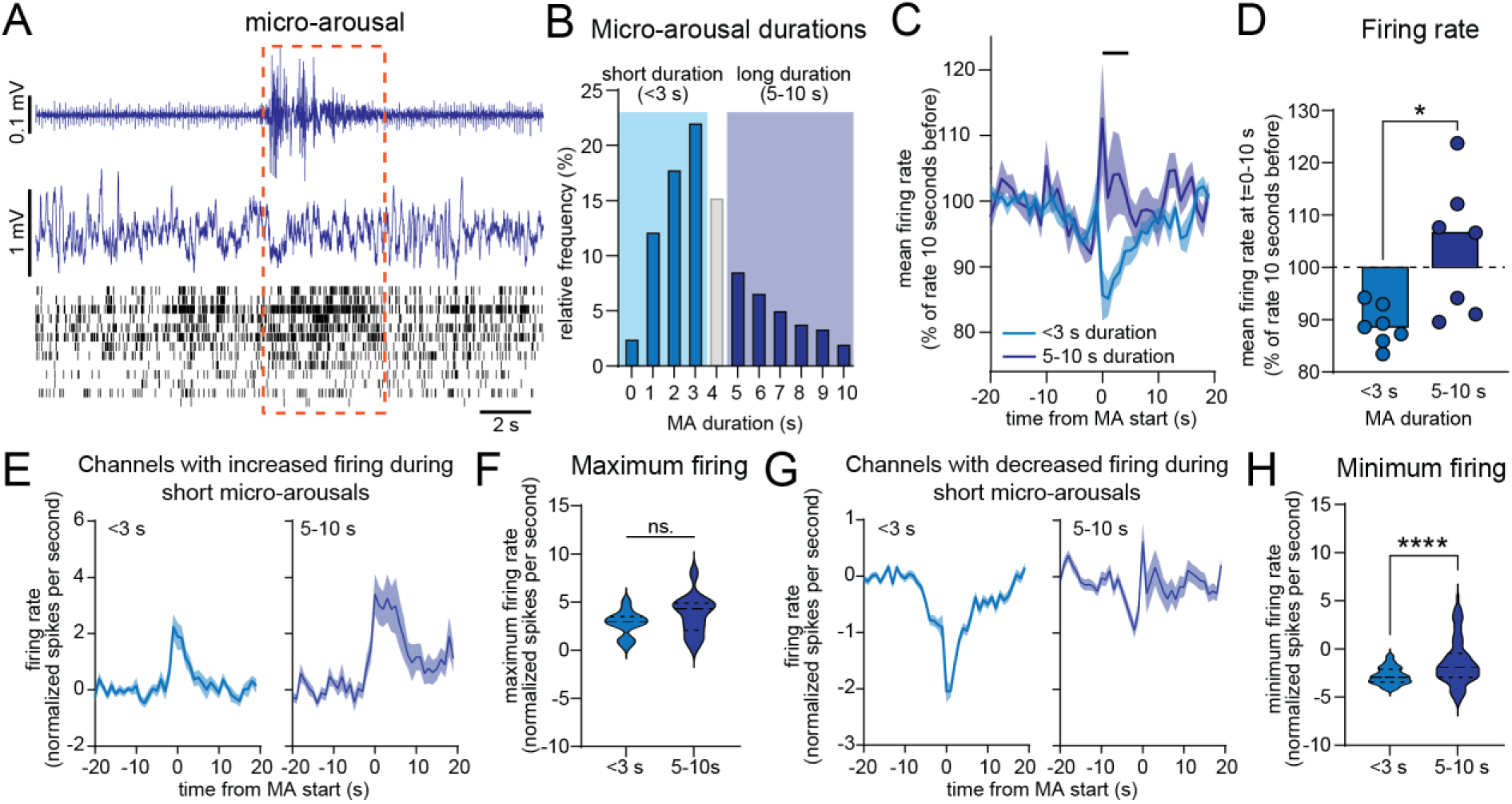
The duration of micro-arousals affects cortical firing. (A) Representative EMG, LFP and multi-unit activity from the motor cortex during a micro-arousal with increased multi-unit activity. (B) Histogram showing the distribution of micro-arousal durations. 3029 micro-arousals in total from 7 mice were analysed. The two tones of blue depict the separation of micro-arousals into groups with durations below 3 s and with durations from 5-10 s. (C) Average number of spikes per second as percent from the firing rate from time-10 to-20. Black line indicates significant values from time 0-3 s (n=7, RM two-way ANOVA with Sidak’s multiple comparison). (D) Average firing rate as percent change from 10 to 20 seconds before the micro-arousal (n = 7, paired t-test, P=0.025). (E) firing rate of channels with increased firing during short (left) and longer (right) micro-arousals calculated as spikes per second normalized to the firing rate and standard deviation during the first 10 seconds of the trace (number of channels=13). (F) Quantification of the maximum firing rate from data in (E) (number of channels=13, paired t-test, dashed lines depict median and interquartile range, P=0.067). (G) firing rate of channels with decreased firing during short (left) and longer (right) micro-arousals calculated as spikes per second normalized to the firing rate and standard deviation during the first 10 seconds of the trace (number of channels=91). (H) Quantification of the maximum firing rate from data in (G) (number of channels=91, paired t-test, dashed lines depict median and interquartile range, P<0.0001). Shaded area around line graphs are ±SEM). MA = micro-arousal.

## Discussion

We here presented an algorithm to automatically detect sleep micro-arousals based on EMG recordings, and used it on data from mice implanted with EMG, EEG and an LFP/multi-unit activity probe placed in the motor cortex. Analysis of the LFP before, during and after micro-arousals showed that all analysed layers (layer 2/3 to 6) of the motor cortex exhibit a uniform decrease in oscillatory power in the SWA (1-4 Hz) range during micro-arousals (Fig. 2). The rate and magnitude of SWA upon micro-arousal to NREM sleep transition was highly sensitive to sleep/wake history, such that mice experiencing low sleep pressure exhibited a long (∼20 s) period of decreased SWA after micro-arousals whereas mice experiencing high sleep pressure exhibited an overshoot in SWA immediately following micro-arousals (Fig. 4). This change could not be explained by a difference in the duration of micro-arousals between the conditions. Importantly, the rate by which SWA decreased as a function of time spent in sleep was steeper for the period immediately following micro-arousals than for the period of NREM sleep from 1 minute before to 1 minute after the micro-arousal. Analysis of single channel firing showed a non-uniform firing pattern where some channels exhibited increased activity immediately prior to micro-arousals while other channels showed decreased firing during micro-arousals (Fig. 5). Finally, longer micro-arousals (5-10 s duration) did not exhibit the drop in firing observed for shorter micro-arousals, which was due to increased firing of low-firing channels.

To our knowledge, this is the first study to identify a relationship between neuronal activity during and around micro-arousals and sleep/wake history. The frequency of micro-arousals has previously been shown to be partly under homeostatic control. Thus, one study reported that 64 hours sleep deprivation in healthy adults resulted in a decrease in micro-arousals during the subsequent recovery sleep^19^, while a study in rats reported that 12 hours of sleep deprivation led to a reduction in the total number of micro-arousals during subsequent recovery sleep^31^. Both studies also reported a significant negative correlation between NREM sleep SWA and the frequency of micro-arousals. A study in mice tested the effect of 4 or 6 hours sleep deprivation on the generation of micro-arousals but only found a robust decrease in micro-arousals after 6 hours of sleep deprivation, while micro-arousals were not significantly affected by 4 hours of sleep deprivation in two of the three mouse strains tested^21^. In addition, this study only found a negative correlation between SWA and the number of micro-arousals in one out of the three mouse strains. Another study reported that sleep deprivation, using methods that are stressful for the mouse, in this case cage shaking, increased both SWA and the number of micro-arousals during subsequent sleep^11^. Thus, the relationship between sleep homeostasis and micro-arousal generation seems to not be straight forward and depend on more factors than the time spent in wakefulness prior to sleep. Our analysis showed a trend toward higher average EMG amplitude during micro-arousals after sleep deprivation, although this effect was not statistically significant. In addition, micro-arousal durations did not differ between low and high sleep pressure states. An interesting next step will be to investigate how the sleep pressure-associated changes in neuronal activity during micro-arousals is affected by experiences during prior wakefulness and to untangle why the number of micro-arousals is negatively correlated with SWA in some studies and not in others.

Our analysis identified a strong correlation between sleep/wake history and SWA in the period immediately after NREM sleep micro-arousals. Surprisingly, the slope of this correlation was steeper than the correlation between sleep/wake history and SWA during a longer period of NREM sleep before and after the micro-arousal, indicating that post-arousal SWA is more sensitive to sleep/wake history than SWA during sustained NREM sleep. The homeostatic increase in NREM SWA in response to high sleep pressure is well-known although its functional significance is still not clear^16–18^. Given our observation that sleep deprived mice exhibited higher SWA levels in association with micro-arousals than during regular NREM sleep, it could be speculated whether micro-arousals play a role in recovery sleep after prolonged wakefulness. Another potential interpretation of the finding that post-arousal SWA was higher in sleep deprived mice could be that it reflects a mechanism that works to terminate the arousal in order to resume sleep. This would explain the increase in SWA as a function of prior time spent in wakefulness, as higher sleep need would impose higher sleep pressure and thereby higher incentive to return to sleep. However, this mechanism would be expected to lead to shorter micro-arousals during high sleep pressure conditions, which we did not observe here. With the tight link between NA and micro-arousals, it is possible that changes in NA dynamics play a role in the increased SWA observed after sleep deprivation. Indeed, NA levels have been shown to decline gradually during sleep and increase during wakefulness^32^, while the amplitude of the NA oscillation is higher after sleep deprivation than during baseline sleep^11,33^. Future studies using combined recordings of NA and neuronal activity after sleep deprivation could provide further insights to this question.

It is also worth noting that while increased SWA after micro-arousals only was observed in sleep deprived mice, all groups exhibited an increase in SWA immediately prior to micro-arousals. What mainly separates the pre-arousal and post-arousal period is that they are on different sites of the rise in NA that promotes the micro-arousal^4^. In other words, the brain has the lowest NA levels during the pre-arousal period and the highest NA levels during the post-arousal period. Evidently, this means that neuronal slow waves before the micro-arousal will happen under very low levels of NA, while slow waves after the micro-arousal will happen while NA release is at its maximum. This difference could impact the downstream processes associated with SWA, such as memory formation. Interestingly, a recent study found that the arousal cycle organizes slow waves in two distinct clusters; a cluster with high amplitude and high degree of synchronization across the scalp, which primarily occurs during high NA levels, and a cluster with smaller and less synchronized slow waves that were more evident during low-arousal periods^34^. NA has been shown to promote long term potentiation (LTP) in the hippocampus^35–37^ and long term depression (LTD) in the dentate gyrus^38^, and reducing the overall levels of NA specifically during sleep leads to impaired memory performance, while enhancing NA during sleep promotes memory formation^39^. Thus, the tightly regulated variability in NA in addition to the hypersynchronous slow firing across the cortical column before and after micro-arousals may provide a window for cognitive processing. This notion is supported by the finding that increased cross-talk between the cortex and the hippocampus takes place during the period leading up to micro-arousals^6^.

Our study suggest that micro-arousals are a heterogeneous phenomenon, and that their properties depend on several factors, including sleep stage, sleep/wake history and duration. For example, REM sleep micro-arousals did not exhibit the increase in SWA prior to the arousal as was seen for NREM arousals, and they did not evoke the same pronounced increase in SWA upon transition back to NREM sleep. In addition, short micro-arousals (<3 seconds) were associated with an overall decrease in cortical neuronal firing, while this was not the case for longer micro-arousals (5-10 seconds). This suggests that arousals of longer duration is more similar to sustained wakefulness, where firing rates are generally higher^30^, while very short bouts of muscle activity is more similar to sleep. Other studies have shown that NA release during NREM sleep not always results in movement and wake-like EEG, but sometimes instead elicits an increase in slow frequencies^4,11^. Although we did not have data for the NA level in this study, it could be speculated whether the very short arousals observed here are closer to the SWA-inducing arousals observed in other studies.

Analysis of single channels revealed a non-heterogeneous pattern, where some channels showed increased activity immediately prior to the first muscle activity, while other channels showed decreased activity during the micro-arousal. Both the channels with increased and decreased firing exhibited higher amplitude changes in sleep deprived mice. Another study recorded neuronal firing in the frontal cortex of rats and found that micro-arousals increased the firing rate of slow-spiking neurons during the subsequent NREM sleep epoch, while epochs of wakefulness had the opposite effect^9^. These results support the notion that micro-arousals are a separate entity from sustained wakefulness, and suggest that micro-arousals may have a role in the homeostatic changes that occur during sleep. A proper identification of single neuronal subtypes would require higher spatial resolution than the 16 channel probes used in this study. An identification of the local neuronal origin of micro-arousals would be valuable in order to explore potential cognitive processes related to micro-arousals and to identify differences between regular and pathological arousals.

There is currently a significant translational gap between the study of micro-arousals in rodents and humans. One of the reasons for this is the discrepancy between how arousals are traditionally defined across human and rodent studies. Thus, scoring of micro-arousals in humans is focused on changes in cortical neuronal activity^40^, while rodent micro-arousal scoring most often is focused on the associated motor response. Additionally, scoring criteria for micro-arousals in rodents is not standardized and therefore often differs between studies, which makes direct comparison of results from different studies difficult. Another reason for why the study of micro-arousals is still at a premature stage is the common misconception, found in both human and rodent studies, that micro-arousals by default is a sign of poor and fragmented sleep. This view is supported by studies showing that acoustically induced arousals during sleep lead to increased sleepiness and reduced performance during the following wake period^41,42^. However, most micro-arousals are internally-generated and occur in the absence of external stimuli^1^. Therefore, externally and internally-initiated micro-arousals most likely don’t have the same implications for sleep continuity and quality. On the other hand, sleep disorders such as sleep apnoea and restless leg syndrome, where repeated awakenings have detrimental effects on the wellbeing of the patient, exhibit a clear cyclical pattern in the generation of arousals, indicating that the infraslow arousal pattern contribute to the disease phenotype^43,44^. Identification of traits that distinguish micro-arousals during healthy sleep and arousals associated with pathological states will be important to detect early signs of pathology, which often manifests as changes in sleep before other clinical signs become evident.

In conclusion, we show that EMG-defined micro-arousals are associated with a uniform drop in LFP power throughout the layers of the motor cortex, and that increased SWA in response to sleep pressure is more pronounced immediately after the micro-arousal than during sustained NREM sleep. In addition, we report that the cortical activity pattern associated with micro-arousals is non-heterogeneous with some channels exhibiting increased firing prior to micro-arousals while other channels exhibit decreased firing and that this firing pattern is amplified under high sleep pressure conditions. The combined set of data shows that internally generated micro-arousals are intimately related to homeostatic regulation of sleep. This finding provides a basis for identifying physiological differences between micro-arousals in healthy, restorative sleep and arousals with detrimental effects on sleep quality and function.

## Methods

All experiments were performed in accordance with the United Kingdom Animal Scientific Procedures Act 1986 under personal and project licenses granted by the United Kingdom Home Office. Ethical approval was provided by the Ethical Review Panel at the University of Oxford. Animal holding and experimentation were located at the Biomedical Sciences Building and the Behavioural Neuroscience Unit, University of Oxford. Chronic electrophysiological recordings from 7 wild-type C57BL/6 mice (125 ± 8 d old at the time of recording) were analysed. This dataset was used in three previous studies^23–25^. In brief, mice were surgically implanted with EEG electrodes for recordings of frontal and occipital regions and a 16-channel silicone probe (NeuroNexus Technologies; A1 × 16-3mm-100-703-Z16, 100 µm spacing between channels) in the motor cortex (+1.1 mm AP, −1.75 mm ML, tilt −15°). A pair of stainless-steel wires were inserted into the nuchal muscle for recording EMG. Following the implantation, mice were housed individually with *ad-libitum* access to food and water for a recovery period of around one week.

## Data collection and signal pre-processing

Sleep recordings were obtained from custom-made Plexiglas cages, which were placed in sound-attenuated and light-controlled Faraday chambers (Campden Instruments) with a 12:12h light:dark cycle and temperature maintained at around 22 °C. Mice had *ad-libitum* access to food and water throughout the recordings. After a three-day acclimatization period where animals habituated to the recording cage and cable, a 24h baseline recording was started at light onset. On the subsequent day at light onset, mice were subjected to 6 hours sleep deprivation by introduction of novel objects to the cage. At the end of the 6 hours, all objects were removed from the cage and the mice were allowed to sleep undisturbed.

The electrophysiological data was acquired using a 128-channel Neurophysiology Recording System (Tucker-Davis Technologies) and the electrophysiological recording software Synapse (Tucker-Davis Technologies), and saved on a local computer. EEG and EMG signals were filtered between 0.1–100 Hz, and stored at a sampling rate of 305 Hz. Local field potentials (LFPs) for each channel were recorded at a sampling rate of 25 kHz and filtered between 300 Hz–5 kHz. Storage of individual spikes was performed whenever the recorded voltage in a laminar channel crossed a manually set threshold to identify putative neuronal spiking (at least two standard deviations above noise level). 46 samples were stored around each spike (0.48 ms before, 1.36 ms after).

EEG, EMG and LFP signals were resampled at a sampling rate of 256 Hz using custom-made MATLAB code (MathWorks, v2017a) and converted into European Data Format as previously described^45^. Spike waveforms were processed using a custom-made MATLAB script to exclude artifactual spikes.

### Scoring of vigilance state

Vigilance states were scored manually by visual inspection of the EMG and frontal and occipital EEG signals at a resolution of 4 s (SleepSign, v3.3.6.1602, Kissei Comtec). Vigilance states were classified as waking (low amplitude, high frequency EEG with EMG activity present), NREM sleep (high amplitude and low frequency EEG with low EMG activity) or REM sleep (low amplitude, high frequency EEG with increased theta activity in the occipital EEG and low EMG).

### Histology

The tips of laminar implants were stained with DiI (Thermo Fisher Scientific) before surgery by immersion of the electrode shank into a 20 mg ml−1 solution (50/50% acetone/methanol) for later histological assessment. After completion of the experiments, micro lesions of selected channels on the laminar probe were performed under terminal pentobarbital anaesthesia using the electroplating device NanoZ (White Matter) applying 10 mA of direct current for 25 s to each respective channel. Immediately following micro lesioning, mice were perfused with 0.1 M PBS (0.9%) followed by 4% paraformaldehyde in PBS for tissue preservation. 50-μm thick coronal sections were cut with a vibratome (Leica VT1000S) and brains were stained with DAPI and mounted on glass slides. Fluorescence microscopy was used to map the depth of the laminar probe implantation and the micro-lesions were used to map the location of specific channels within the cortical layers.

### Micro-arousal detection

In order to make the analysis of neuronal activity around micro-arousals as un-biased as possible, the detection of micro-arousals was based solely on the EMG signal and not the EEG signals. Furthermore, the analysis was mainly focused on micro-arousals during NREM sleep, so micro-arousals following REM sleep were not included, except in a version of the algorithm used specifically to detect micro-arousals after REM sleep. A variety of methods are used for vigilance state scoring and range from fully-or semi-automatic scoring (see e.g. ^46–49)^ to manual scoring where vigilance states are assigned by visual inspection. Therefore, the goal was not to develop an algorithm for general sleep scoring, but one that could be used on already scored data to extract information about micro-arousals specifically.

As a first step, intruding periods scored as wakefulness or REM sleep during NREM sleep were re-scored as NREM sleep if the duration of the period was below 10 seconds. This was done to remove existing manually scored micro-arousals. The EMG signal was then filtered and rectified using the moving standard deviation, the moving mean and a 10000^th^ order median filter. Peaks in the EMG signal was detected if the signal crossed a threshold of 2 * the standard deviation. In the subsequent steps, EMG peaks were excluded if they did not occur within periods of NREM sleep, and EMG peaks that were less than 10 seconds apart were combined as one event. Finally, EMG peaks with a duration above 5 or 10 seconds, depending on the analysis in question, were excluded. As a result, a vector with the beginning of each micro-arousal and a vector with each micro-arousal duration is generated and stored.

The detection algorithm was validated by visual inspection of the EMG and EEG signal and the automatically scored micro-arousals for each mouse, as well as calculation of the average muscle activity during automatically detected micro-arousals compared to micro-arousals scored by manual scoring (figure 1).

For detection of micro-arousals following REM sleep, the algorithm was adapted to only include micro-arousals that occurred within 4 seconds after REM sleep termination. Because EMG activity after REM sleep showed more variability across mice, thresholds for detection of EMG peaks were decided for each mouse after visual inspection of the detected micro-arousals.

## Data analysis

All data analysis was performed in MATLAB (R2024A). Analysis of specific cortical layers were performed on a single channel confirmed to be within the layer of interest from post-mortem histological analysis.

Prior to spectral analysis, data traces were filtered using a 20^th^ order bandpass Butterworth IIR filter with a lower cut-off at 1 Hz and a higher cut-off at 30 Hz. The lower cut-off of 1 Hz was chosen to exclude potential low frequency artefacts caused by movement during micro-arousals.

### Spectrograms

Spectrograms were generated using the “spectrogram” function with a window of 500 ms and an overlap of 0. Mean power within specific power bands was calculated from the spectrogram by averaging the power within the frequencies of interest. Power spectral density (PSD) plots were calculated using pwelch with a frequency resolution of 0.5 Hz, zero overlap and a Hanning window. PSD plots from before, during and after a micro-arousal were calculated 30-50 seconds before the micro-arousal, 0-2 seconds after the beginning of the micro-arousal and 5-10 seconds after the beginning of the micro-arousal, respectively.

### Correlation between SWA and sleep/wake history

For plotting the correlation between sleep/wake history and post-micro-arousal SWA, all micro-arousals with a duration below 5 seconds occurring after 6-hour sleep deprivation were detected and used in the analysis (sleep deprivation started at lights on so micro-arousals during the 6 last hours of lights on and 12 hours during lights off were included). SWA 3-8 seconds after the beginning of each micro-arousal was first calculated and normalized to the level of SWA during the light phase of the baseline recording from the same mouse. Then, the percent time the mouse had spent in NREM sleep during the 6 hours before each micro-arousal was calculated, and the normalized post-micro-arousal SWA was plotted against the percent time spent in NREM sleep. To plot the relationship between SWA during regular NREM sleep and sleep/wake history, SWA during a period from 1 minute before to 1 minute after each micro-arousal was calculated and normalized to SWA during the light phase of the baseline recording. This value was plotted against the percent time the mouse had spent in NREM sleep prior to the micro-arousal.

### Calculation of average firing rates

The average firing rate from multi-unit recordings was calculated as the combined number of spikes from all channels per second. The firing rates were first calculated for each micro-arousal and then averaged across micro-arousals. For comparison between mice, the averaged firing rate was normalized as the percentage change from the first 10 seconds of the trace (10 to 20 seconds before the micro-arousal).

Analysis of single-channels was performed on the averaged firing rates. To compare firing rates across animals and sleep pressure conditions, the firing rate during micro-arousals under the high sleep pressure condition was used to calculate the standard deviation of the firing rate and the mean of the first 10 seconds for each mouse. These values were subsequently used to normalize the firing rates for all conditions by dividing with the standard deviation and subtracting the mean.

For comparison of single channels across sleep pressure conditions, channels with decreased firing were detected by collecting all channels during the high sleep pressure condition where the minimum value during a period from 5 seconds before to 5 seconds after the micro-arousal was below the mean firing rate 5 to 15 seconds before the micro-arousal minus 5 times the standard deviation. Likewise, channels with increased firing during the high sleep pressure condition were detected by collecting all channels where the maximum value during a period from 5 seconds before to 5 seconds after the micro-arousal was above the mean firing rate 5 to 15 seconds before the micro-arousal plus 5 times the standard deviation. The firing rate of channels detected as increased or decreased during the high sleep pressure condition were then compared to the firing rate of the same channels during low and medium sleep pressure conditions.

Comparison of single channels between different micro-arousal durations followed the same method, except normalization values were calculated for micro-arousals with a duration below 3 seconds, and all micro-arousals during the 24-hour baseline period were used. Furthermore, the criteria for selection of channels with increased or decreased firing was based on firing above or below 2 times the standard +/-the mean firing rate 5 to 15 seconds before the micro-arousal was used. This was to ensure a sufficient inclusion of channels as fewer channels showed increased firing during short micro-arousals.

### Calculation of average rectified EMG

Average rectified EMG traces were calculated by taking the mean of the filtered and rectified EMG trace time-locked to each micro-arousal.

## Statistics

Statistics were done in GraphPad Prism (version 10.3.1). P-values less than 0.05 (after correction for multiple comparison except the comparison of power density per frequency bins) were considered significant. For comparison of the means of two group where the groups consisted of the same mice under different conditions or time bins, a two tailed paired t-test were used. For comparison between the means of three or more groups, RM one-way ANOVA with Geisser-Greenhouse correction and Tukey’s multiple comparison was used. For comparison of power densities across frequency bins RM two-way ANOVA with matched values stacked into sub columns was used. Pearson correlation was used to calculate correlation coefficients and slopes of the correlations. ANCOVA was used to test for significant difference between linear regression lines.

## Supplementary figures

**Figure S1.**
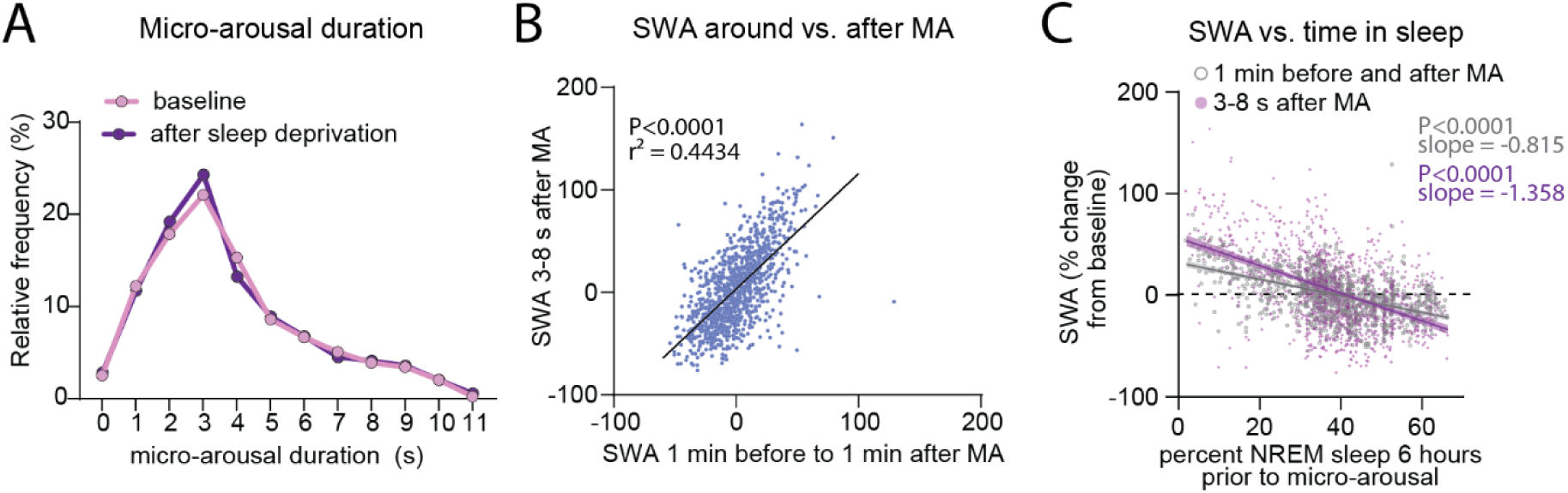
(A) Histogram showing the relative frequency distribution of micro-arousal durations during baseline and after sleep deprivation (n=7). (B) SWA measured 1 minute before to 1 minute after each micro-arousal correlated to SWA 3-8 seconds after the same micro-arousal (Pearson correlation, 1162 micro-arousals from 7 mice during 18 hours after 6 hours sleep deprivation included, simple linear regression). (C) Percent time spent in NREM sleep during a period 6 hours prior to each micro-arousal correlated with the amplitude of SWA around (1 minute before to 1 minute after, shown in grey) or after (3-8 seconds after, shown in purple) the same micro-arousal. SWA amplitude is shown as percent of mean NREM SWA during the first 6 hours of lights on. 1162 micro-arousals from 7 mice during 18 hours after 6 hours sleep deprivation included. The lines show the linear regression and the shaded area depict 95% confidence intervals. Comparison of the slopes with linear regression showed a significant difference (Pearson correlation, P<0.0001, ANCOVA). MA = micro-arousal, SWA = SWA.

**Figure S2.**
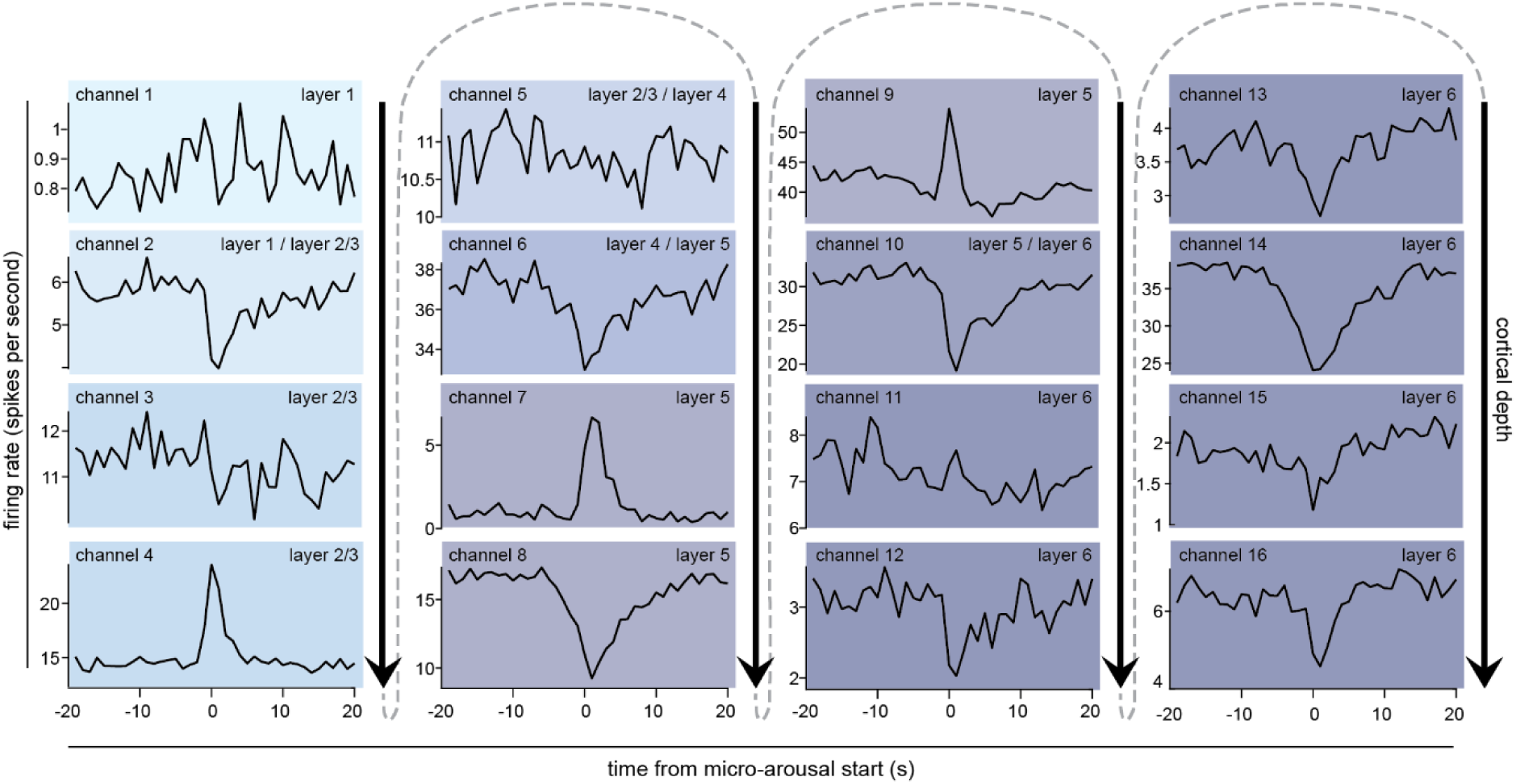
Example firing rates around micro-arousals from one mouse. Each plot corresponds to the number of spikes per second for a single LFP channel. Colour coding shows increasing cortical depth. Note that channels exhibiting decreased and increased firing are spread across the cortical column and are not separated into specific layers.

